# Genetic mapping of multiple metabolic traits identifies novel genes for adiposity, lipids and insulin secretory capacity in outbred rats

**DOI:** 10.1101/2022.03.14.484225

**Authors:** Thu H Le, Wesley L Crouse, Gregory R Keele, Katie Holl, Osborne Seshie, Michael Tschannen, Ann Craddock, Swapan K. Das, Bailey McDonald, Neeraj K Sharma, Chia-Chi Chuang Key, Gregory Hawkins, William Valdar, Richard Mott, Leah C Solberg Woods

## Abstract

Despite the successes of human genome-wide association studies, the causal genes underlying most metabolic traits remain unclear. We used outbred heterogeneous stock (HS) rats, coupled with expression data and mediation analysis, to identify quantitative trait loci (QTLs) and candidate gene mediators for adiposity, glucose tolerance, serum lipids and other metabolic traits. Physiological traits were measured in 1519 male HS rats, with liver and adipose transcriptomes measured in over 410 rats. Genotypes were imputed from low coverage whole genome sequence. Linear mixed models were used to detect physiological and expression QTLs (pQTLs and eQTLs, respectively), employing both SNP- and haplotype-based models for pQTL mapping. Genes with cis-eQTLs that overlapped pQTLs were assessed as causal candidates through mediation analysis. We identified 15 SNP-based pQTLs and 19 haplotype-based pQTLs, of which 11 were in common. Using mediation, we identified the following genes as candidate mediators of pQTLs: *Grk5* for a fat pad weight pQTL on Chr1, *Krtcap3* for fat pad weight and serum lipids pQTLs on Chr6, *Ilrun* for a fat pad weight pQTL on Chr20 and *Rfx6* for a whole pancreatic insulin content pQTL on Chr20. Furthermore, we verified *Grk5* and *Ktrcap3* using gene knock-down/out models, thereby shedding light on novel regulators of obesity.

## INTRODUCTION

Obesity and overweight are major risk factors for poor metabolic health, including dyslipidemia, cardiovascular disease and diabetes[1]. Both genetic and environmental factors contribute to obesity and related metabolic traits, with up to 70% of the population variance attributable to genetics[2]. Although human genome-wide association studies (GWAS) have associated many genomic regions with lipid levels[3], fasting glucose and insulin[4,5], diabetes[6], body mass index (BMI)[7], and waist-hip-ratio (WHR) [8,9], causal genes in most of these loci remain unknown. This is due, in part, to the challenge of collecting gene expression and other -omic phenotypes from relevant human tissues.

Animal models of obesity have several advantages over human studies; environmental and dietary factors are controllable, and -omic data such as transcriptomics can be collected from relevant tissues under controlled conditions. In particular, the heterogeneous stock (HS) rats are an established model for the genetic study of obesity[10,11]. The HS are descended from eight fully sequenced inbred founder strains [12,13]. The HS genomes are highly heterozygous, and each chromosome is a fine-grained mosaic of the founder haplotypes, suitable for high resolution genetic mapping [14,15].

Here, we use the HS rat model to dissect the genetic architecture of metabolic and adiposity-related phenotypes. This work significantly extends our previous study [10,11] in which we mapped physiological QTLs (pQTL) for adiposity traits. First, we use many more animals, assess additional metabolic traits and measure gene expression in two metabolically active tissues. Second, instead of genotyping the animals at a limited set of SNPs with an array, we genotype at a comprehensive genome-wide set of sites using low coverage whole genome sequencing (LC-WGS) followed by imputation. Third, we use mediation analysis to identify candidate causal genes that act in a tissue-specific manner, and show that many pQTL are likely regulated by multiple genes. Finally, we validate two novel genes, *Krtcap3* in liver and *Grk5* in adipose, using in vivo knock-out and cell gene silencing models, respectively.

## RESEARCH DESIGN AND METHODS

### Animals and phenotypes

The outbred NIH HS rat colony was initiated in 1984 from eight inbred founder strains: ACI/N, BN/SsN, BUF/N, F344/N, M520/N, MR/N, WN/N, WKY/N, and maintained to minimize inbreeding[12]. Rats used in this study were maintained at The Medical College of Wisconsin (NMcwi:HS; RGD_2314009) and housed as described in **Supplementary Methods**. 1519 male rats were phenotyped using the protocol in [10]. Briefly, at age 16 weeks, we performed an intra-peritoneal glucose tolerance test (IPGTT) on each rat. After an overnight fast, we measured body weight (BW) and basal fasting glucose (Gluc0) and insulin (Ins0). We then injected rats with 1mg/kg glucose, collecting blood glucose and serum for insulin at 15, 30, 60, 90 minutes post injection. We calculated glucose area under the curve (GluAUC), insulin area under the curve (InsAUC) and quantitative insulin sensitive check (QUICKI), a measure of insulin sensitivity [16]. At 17 weeks of age, a sub-set of 1144 rats were euthanised after an overnight fast, after which we measured body weight (sacBW) and body length with and without tail (BLTail and BLNoTail, respectively). We used an enzymatic detection method to measure fasting serum total cholesterol (CHOL) and triglycerides (TRIG) from trunk blood. We dissected and weighed the heart, pancreas, retroperitoneal (RetroFat) and epididymal visceral fat pads (EpiFat), and calculated Body weight gain (BWGain) as the difference between BW at sacrifice and BW at IPGTT (ie.sacBW-BW). Whole pancreas insulin content (WPIC) and serum insulin during the IPGTT were also measured using an ELISA kit (Alpco Diagnostics, Salem, NH). Liver and adipose tissues were snap-frozen for expression analyses. In order to collect urinary protein levels for another study[17], we euthanised 375 rats at 24 weeks, and weighed left kidney (LftKid), right kidney (RigKid), and both kidneys (TotKid). All protocols were approved by the Institutional animal care and use committee at MCW. Phenotype data have been deposited in the Rat Genome Database (www.rgd.mcw.edu).

Prior to genetic analysis, tissue weights were adjusted to account for body weight by dividing by sacBW. Each phenotype was transformed to approximate normality using either the log, rank-inverse normal transformation (RINT) or a more conservative variant of RINT (TRINT), as detailed in **Supplemental Methods** and **Supplemental Table 1**.

### RNA-Seq

RNA from liver and adipose tissue was extracted from subsets of 430 and 415 rats, respectively, selected to maximize genetic diversity (e.g., no more than 1 or 2 rats per family), while encompassing the phenotypic spectrum of fat pad weights. Library created using Illumina kits were run on an Illumina HiSeq 2500 to obtain 37bp paired-end reads for liver and 75bp single-end reads for adipose tissue. We used STAR [18] to align reads to the reference Rn6.0 and DESeq2 [19] to compute gene level expression counts. We excluded very lowly expressed genes with average reads per sample < 1. The expression of 18,358 genes from adipose and 16,796 genes from liver were then RINT-transformed for analyses.

### Low coverage whole genome sequencing and genotype imputation

DNA from the HS rats was sequenced at BGI using 143bp paired-end Illumina reads with mean coverage of 0.24x, followed by imputation using STITCH [20]. Pre-existing Illumina high-coverage (24-28X) sequence from the eight founder strains [13] provided a haplotype reference panel. After quality control (imputation info score > 0.4 and Hardy-Weinberg Equilibrium p-value >10^−6^), we retained 4,865,047 imputed SNPs. For genetic mapping, we used Plink [21] to remove SNPs in high linkage disequilibrium (LD) (r^2^ > 0.95), leaving 125,611 SNPs for genetic mapping.

### Statistical analyses

#### Founder haplotype dosages and kinship matrices

Founder haplotype dosages were computed at each of the ~126K tagging SNPs using STITCH [20]. To account for the unequal relatedness between HS rats, we estimated a kinship matrix in two ways. The first was a SNP-based additive kinship matrix, ***K***_*SNP*_, computed from the tagging SNPs, which was used to estimate heritability and genetic correlations for the physiological traits, and in genetic mapping of the expression traits. The second was a haplotype-based kinship matrix, ***K***_*HAP*_, used for genetic mapping of physiological traits. This was estimated in R/qtl2 [22] using the afformentioned founder haplotype dosage information from either all chromosomes (used for SNP-based mapping) or all but one chromosome (used for haplotype-based mapping; see **Supplemental Methods**).

#### Genetic mapping and analyses of physiological traits

The normalised physiological phenotypes and 126K tagging SNPs were used for all genetic analyses. We conducted both SNP-based and haplotype-based genetic mapping to identify physiological QTLs (pQTLs).

We mapped SNP-based pQTLs using imputed SNP dosages from STITCH with the kinship matrix ***K***_*HAP*_, via the miqtl R package [10]. The mixed model for SNP-based pQTL mapping at the SNP indexed by position *s* is

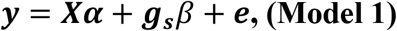

where ***y*** is the vector of normalized phenotypes, ***X*** is a design matrix of fixed effect covariates, including coat color, euthanasia order, tissue harvest order, and technician (**Supplemental Table 1**), ***α*** is the vector of regression coefficients for these fixed effects, and ***g***_***s***_ is the vector of SNP dosages with regression coefficient *β*. The error vector ***e*** has variance-covariance matrix 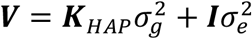, where 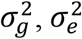 are respectively the additive genetic and environmental variance components for the trait. Pearson correlations between physiological traits were estimated in R. We tested association between phenotype and SNP using generalized least squares by pre-multiplying the above equation by the inverse of the matrix square root of ***V***. We estimated genome-wide significance thresholds by parametric bootstrap, performing genome scans on 500 simulations from a fitted null model (i.e., no QTL) [23,24]. Confidence intervals on pQTL position were defined using LD drop between the peak and neighbouring SNPs, using a threshold ***R***^2^ of 0.5.

**Table 1 -.**
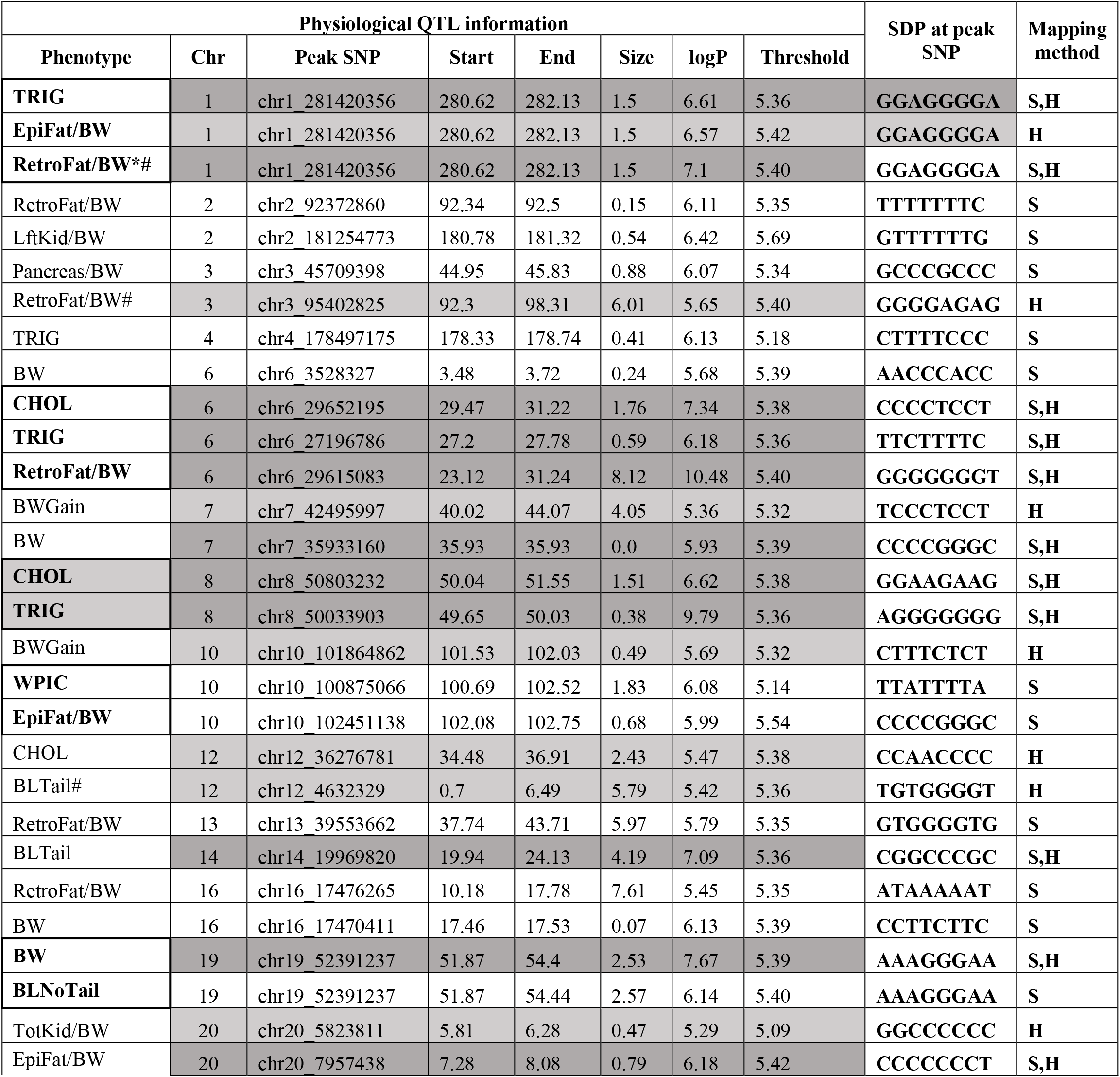
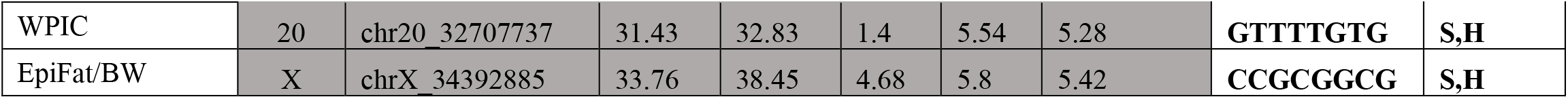
**Physiological QTLs (pQTLs) detected in up to 1519 male HS rats. Each row shows one pQTL. The strain distribution pattern (SDP) at the peak SNP of the eight founders uses the following order: ACI-BN-BUF-F344-M520-MR-WN-WKY. In terms of mapping method, pQTL detected by SNP-based only (S) are in white background, Haplotype-based only (H) are in light grey background or both (S,H) methods are in dark grey background. Five pQTLs affecting multiple physiological traits were grouped together in the first column and marked in bold. pQTLs that replicate those reported in**[10] **or** [11] **are indicated by * and # respectively**.

Haplotype-based mixed-model genetic mapping was performed using R/qtl2 with the haplotype dosages, together with the kinship ***K***_*HAP*_, with equation

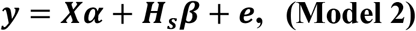

where ***H***_***s***_ is the matrix of haplotype dosages at site *s*. The genome-wide significance threshold was estimated using permutation and pQTL positional confidence intervals were defined by extending the location of the peak SNP to both sides until all the SNPs within that region had LOD score higher than the peak LOD score minus 1.5 (i.e., 1.5 LOD drop). We converted LOD scores to logP units given that under the null hypothesis 2 log_e_10 * LOD is distributed as a 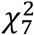 random variable.

Heritability and genetic correlations were estimated using GCTA [25] in a bi-variate linear mixed models similar to Models 1 but with the SNP effects excluded. Heritability was estimated as 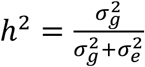.

#### Estimation of founder haplotype effect at physiological QTLs

We refined our estimates of founder haplotype effects at each detected QTL using Diploffect [26] to determine which founders carried non-zero haplotype effects on physiological phenotypes (pQTLs) and gene expression (eQTLs), and whether the QTL is potentially multi-allelic.

#### Genetic mapping of liver and adipose expression levels

We used Matrix eQTL [27] to map eQTL with the same genotypes and kinship matrix ***K***_SNP_ restricted to those rats with expression data. We tested normalized gene expression levels for association with SNP dosages in a linear mixed model, as with SNP-based pQTL mapping, except the choice of covariates (**Supplement Table 1**). For adipose tissue, covariates include one proxy for tissue composition constructed from 27 highly variable and mainly muscle-expressed genes, one nerve marker gene (MPZ), batch and the first 10 genotype principal components (PCs), as described in **Supplemental Methods**. For liver genes, covariates included batch, RNA integrity number (RIN), the first PC from expression data and the first 10 PCs from genotypes. We focused on cis-eQTLs in order to identify candidate mediators of pQTLs. A cis-eQTL for a gene was recorded if at least two associated SNPs with logP>5 mapped within 1 Mb of that gene’s location.

#### Mediation analysis

We conducted mediation analysis to assess whether a gene with a cis-eQTL containing at least two associated SNP that falls within the interval of a pQTL acts as an intermediate for the physiological trait using the methodology in [10,28]. Briefly, consider gene *z* with a cis-eQTL mapping within the interval of a pQTL. One potential explanation is that the eQTL and pQTL are driven by the same causal variant, and that the action of the pQTL on the phenotype is partially or fully mediated through the transcript level of gene *z*. The evidence for this is evaluated by comparisons among three models:

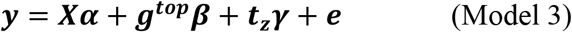

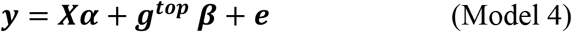

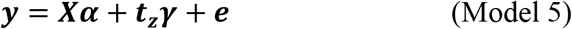

where ***y*** is the normalized physiological phenotype, ***t***_***z***_ is the normalised expression for gene *z*, and ***g***^***top***^ is the SNP genotype dosage for the top SNP within the pQTL. Positive evidence for mediation is reported when Model 3 is significantly more predictive of the phenotype than Model 4. Because this test is applied to all *n*_*t*_ genes with cis-eQTL within a given pQTL, the *n*_*t*_ p-values are subject to multiple test correction using the Benjamini-Hochberg false discovery rate (FDR), with significant mediators defined as genes with q-values < 0.1. For significant mediators, full mediation is implied when Model 3 fails to be significantly more predictive than Model 5 (nominal p-value < 0.05); otherwise, mediation is considered partial.

#### Transduction of 3T3-L1 cells to knock down Grk5 expression

We grew 3T3-L1 cells, a mouse preadipocyte cell line, under standard culture conditions [29]. The *Grk5* gene was silenced by infecting 3T3-L1 preadipocytes with lentiviral particles to deliver mouse gene-specific shRNA expression vectors (sc-39043-V; Santa Cruz Biotechnology, Santa Cruz, CA) in the presence of polybrene (sc-134220 [8 microgram/mL]; Santa Cruz Biotechnology) according to the manufacturer’s protocol. Control shRNA, lentiviral Particles-A (sc-108080; Santa Cruz Biotechnology) were used as a negative control. Cells successfully transduced and stably expressing shRNA were selected using 2 microgram/mL puromycin (A1113803; Gibco, Thermo Fisher Scientific). The RNA of control and puromycin-selected *Grk5* knock-down 3T3-L1 (n=3 per genotype) cells were isolated and reverse-transcribed into cDNA for real-time PCR quantification of *Grk5* (TaqMan primer #Mm00517039_m1) and *Gapdh* (an endogenous control, TaqMan primer # Mm99999915_g1) to determine the extent of *Grk5* knock-down.

#### Adipocyte differentiation and TRIG measurement

The 3T3-L1 preadipocytes were differentiated into adipocytes for 7 days as described previously [30,31]. Cellular TRIG was measured from nonsaponified samples after solubilization in Triton-X-100, using a commercially available kit (Wako L-Type Triglyceride M test), and normalized to protein content (Pierce™ BCA Protein Assay Kit) [31].

## RESULTS

### Metabolic traits are highly correlated and show strong heritability

SNP-based heritabilities, genetic correlations and phenotype correlations of the normalized physiological traits are plotted in **Figure 1** and tabulated in **Supplementary Tables 3**. Parameters for total tissue weights (prior to dividing by body weight) are shown in **Figure 1**, and tissue weight fractions are in **Supplementary Figure 1**. Heritabilities ranged between 0.18 and 0.60 and are typical for rat studies of these phenotypes [10]. Heritability was highest for BW and fat pad weights (RetroFat and EpiFat) and lowest for BWGain and BLNoTail (0.54 to 0.6 vs 0.18 to 0.22). Phenotypic (*R*_*p*_) and genotypic (*R*_*g*_) correlations between traits tracked each other closely but on average *R*_*g*_ is 19% larger than *R*_*p*._ Fat pad weight traits, body length (BLTail and BLNoTail) and BW were significantly positively correlated (0.4 < *R*_*g*_ < 0.95). These measures were more strongly correlated with insulin traits (0.25 < *R*_*g*_ < 0.73) than with glucose traits (−0.03 < *R*_*g*_ < 0.28). Fat pad weights were also significantly positively correlated with other metabolic traits: *R*_*g*_ ~ 0.45 with TRIG and *R*_*g*_ ~ 0.35 with CHOL.

**Figure 1 -.**
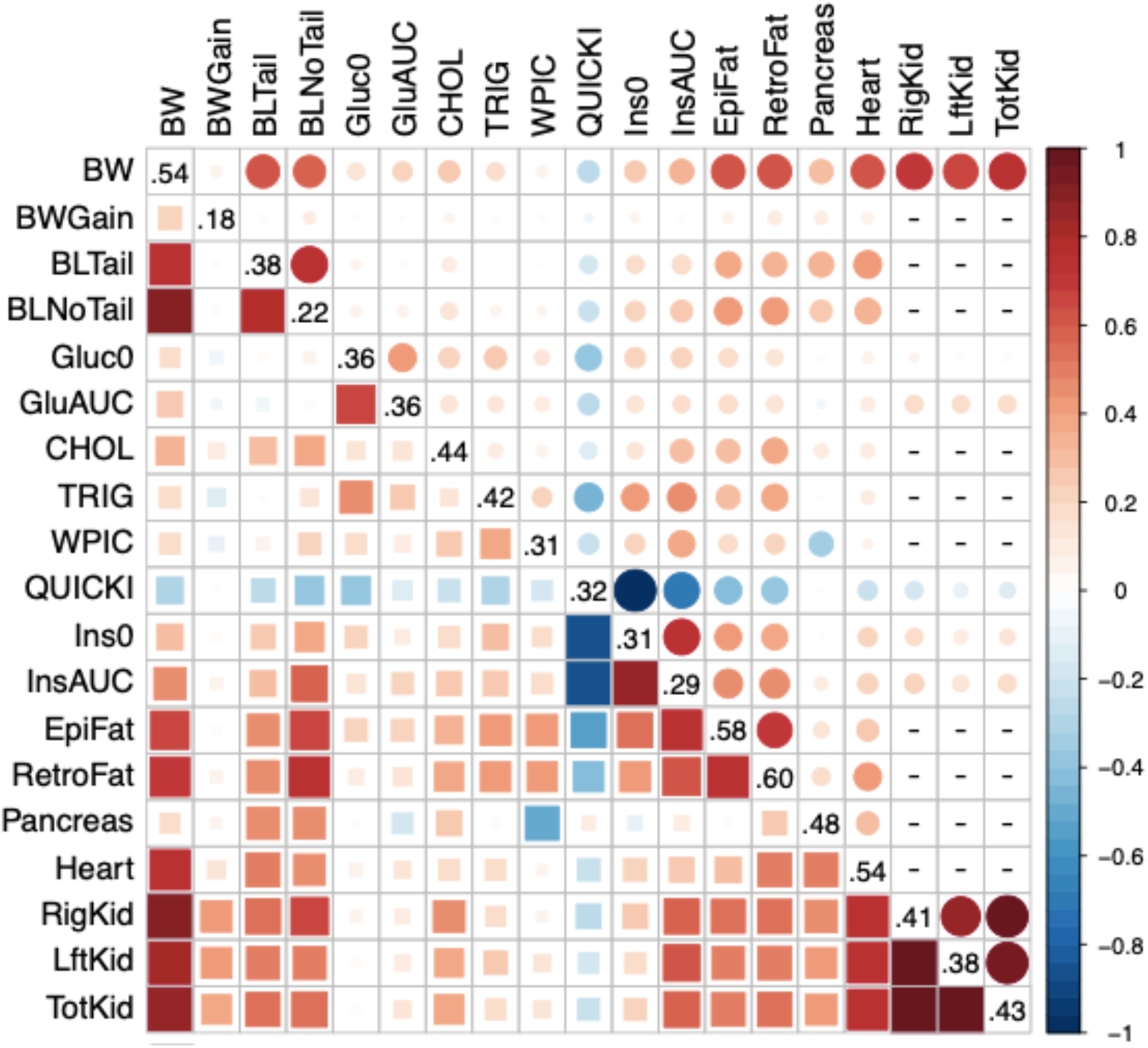
Heritabilities, genetic and phenotypic correlations for physiological traits in up to 1519 male HS rats. The main diagonal contains the heritabilities, the upper triangle the phenotypic correlations and the lower triangle the genetic correlations. Dashes indicate missing data for phenotypic correlations, where traits were measured on different animals.

### Physiological QTLs

All pQTL exceeding the 10% genome-wide significance thresholds and the strain distribution patterns (SDP) at the top SNPs of each pQTL are shown in **Table 1** and **Figure 2**. We mapped 15 SNP-based pQTL and 19 haplotype-based pQTL, of which 11 overlap. pQTL mapping interval widths range between 0.59 and 8.12 Mb for SNP-based pQTLs and between 0.07 and 7.61 Mb for haplotype-based pQTLs. Several pQTLs detected by both SNP- and haplotype-based mapping and, based on haplotype plots (**Supplementary Figure 2**), are likely pleiotropic: pQTLs on Chr1:180Mb associate with TRIG, EpiFat/BW and RetroFat/BW; pQTLs on Chr6:27Mb associate with CHOL, TRIG and RetroFat/BW; pQTLs on Chr8:50Mb associate with CHOL and TRIG; pQTLs on Chr10:101Mb associate with WPIC and EpiFat/BW; and pQTLs on Chr19:52Mb associate with BW, BLNoTail and EpiFat. We also identified a pQTL on Chr16 that associate with both BW and RetroFat/BW using only haplotype-based mapping. Other pQTLs mapped to overlapping regions but are unlikely to be pleiotropic based on having differing haplotype effects (**Supplementary Figure 2**): Chr10: 101Mb for BWGain overlaps the Chr10:101Mb QTL for WPIC and EpiFat/BW; Chr20:6Mb maps TotKid/BW and EpiFat. Fat pad weights that were not divided by sacBW mapped to generally the same regions as those divided by sacBW as shown in **Supplementary Table 4**. Several glucose and insulin measures lacked genome-wide significant pQTL (**Supplementary Figure 3**) despite strong heritability. We identified a likely missassembly in the Rn6.0 reference for Heart Weight on Chr5, where two pQTL separated by 50Mb were in strong LD (shown as diamonds in **Figure 2**). On reference Rn3.4 these loci are linked, and the pQTLs represent a single region.

**Figure 2 -.**
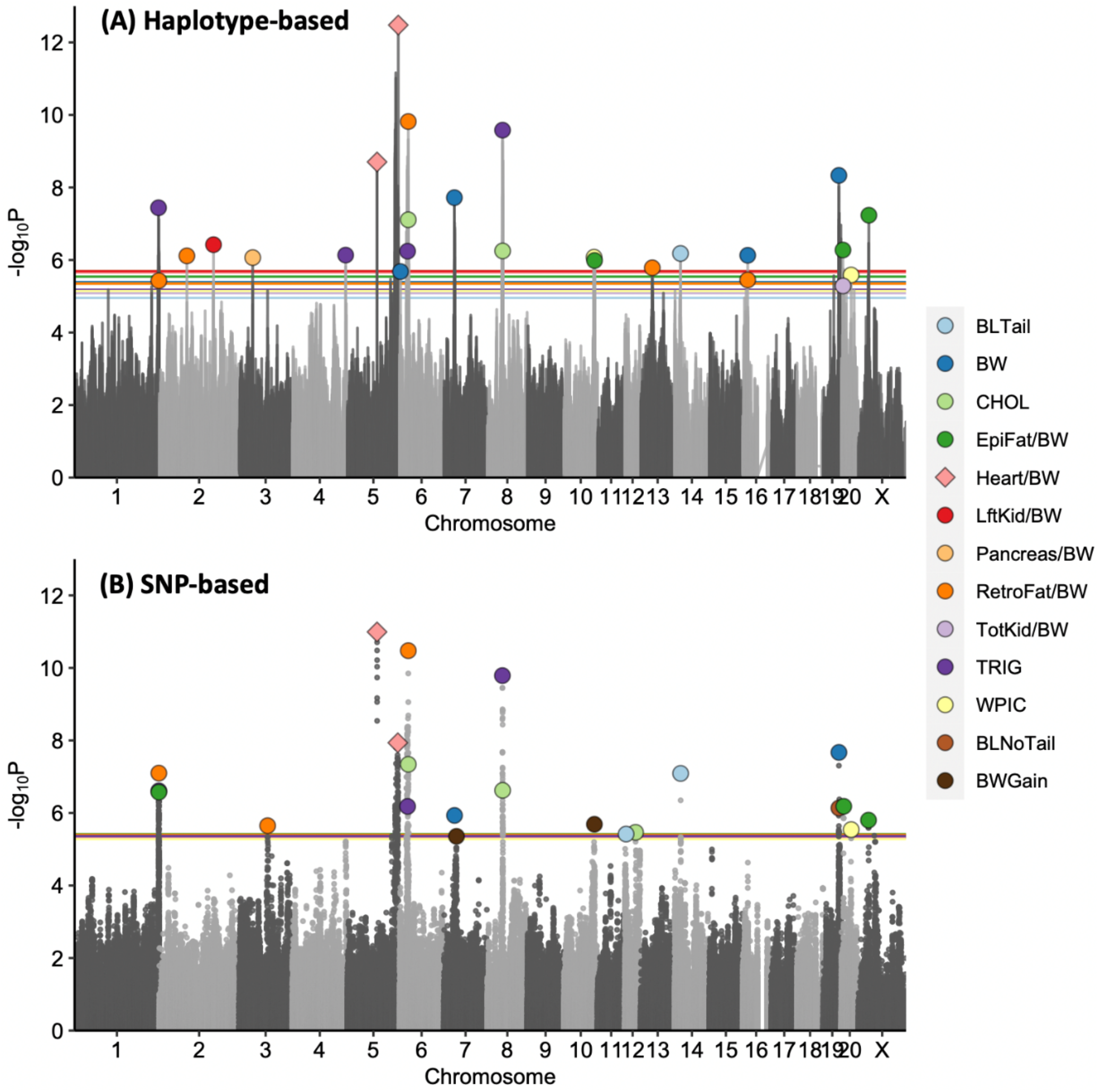
Porcupine plot of pQTLs detected as genome-wide significant using (A) haplotype-based or (B) SNP-based mixed models. The x-axis shows genomic position and y-axis the significance as negative log 10 of the p-value of the test of association. Traits are color-coded as indicated in legend. Thresholds are shown as horizontal lines and indicate the genome-wide 10% thresholds as determined by permutation in haplotype and bootstraps in SNP-based mapping. Haplotype association thresholds vary between traits whereas those for SNPs are almost the same for all traits. Two pQTLs for Heart weight indicated by diamonds are likely the same locus, probability due to a missassembly in Rn6.0.

**Figure 3 -.**
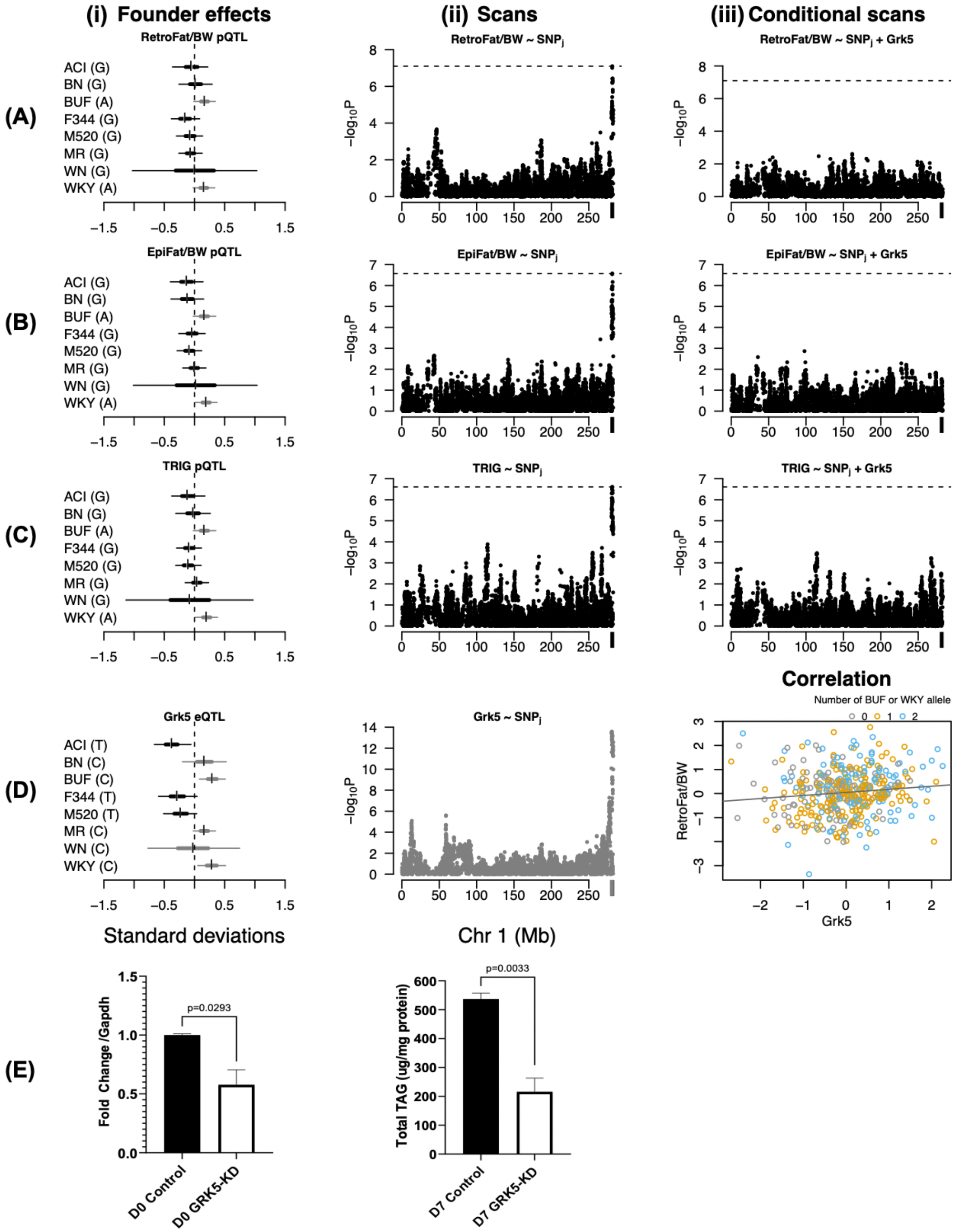
Adipose expression of *Grk5* is a full mediator for RetroFat/BW and EpiFat/BW; and a partial mediator for Triglycerides (TRIG). Each row A-C shows mediation analysis for the phenotypes RetroFat/BW, EpiFat/BW and TRIG respectively. Within each row, left to right, are shown (i) box-whisker plots of the estimated haplotype effects with uncertainties at the peak SNP for the phenotype; (ii) the scan across chromosome 1 for SNP-based mixed model for the trait; and (iii) the corresponding scan after including the expression of *Grk5* as a covariate. Row D shows the expression QTL analysis for *Grk5*, in which the right-most panel is a scatter plot of RetroFat/BW vs *Grk5* expression in adipose tissue across 415 rats, color-coded by genotype at the peak SNP of the pQTL for RetroFat/BW, which is private to the founder WKY and also indicates the estimated number of WKY haplotypes. Row E shows that knock-down of *Grk5* decreases total TRIG accumulation in a 3T3L1 adipocyte cell line.

### Cis-Expression QTLs

We analysed 16,796 and 18,358 transcripts with detectable expression levels in liver and adipose, respectively. Using SNP-based mixed models we mapped 2,267 liver and 2,226 adipose cis-eQTLs with Benjamini-Hochberg false discovery rates (FDR) of 0.00024 for liver and 0.00028 for adipose (**Supplement Table 1, 5 and 6**).

### Mediation analysis to identify candidate genes

We performed mediation analysis to identify candidate causal genes with cis-eQTLs in either tissue that overlapped a SNP-based pQTL. The number of genes with overlapping cis-eQTLs in each pQTL ranged from 1 to 15 in liver and from 1 to 14 in adipose (average number of 5 genes), cumulatively representing 238 candidate genes with cis-eQTLs within pQTLs (**Supplemental Table 7**). We tested if the expression of the gene fully or partially absorbed the effect of the pQTL. We identified four full mediators (three in liver and one in adipose) and 41 partial mediators (17 in liver and 24 in adipose), listed in **Table 2**.

**Table 2 -.**
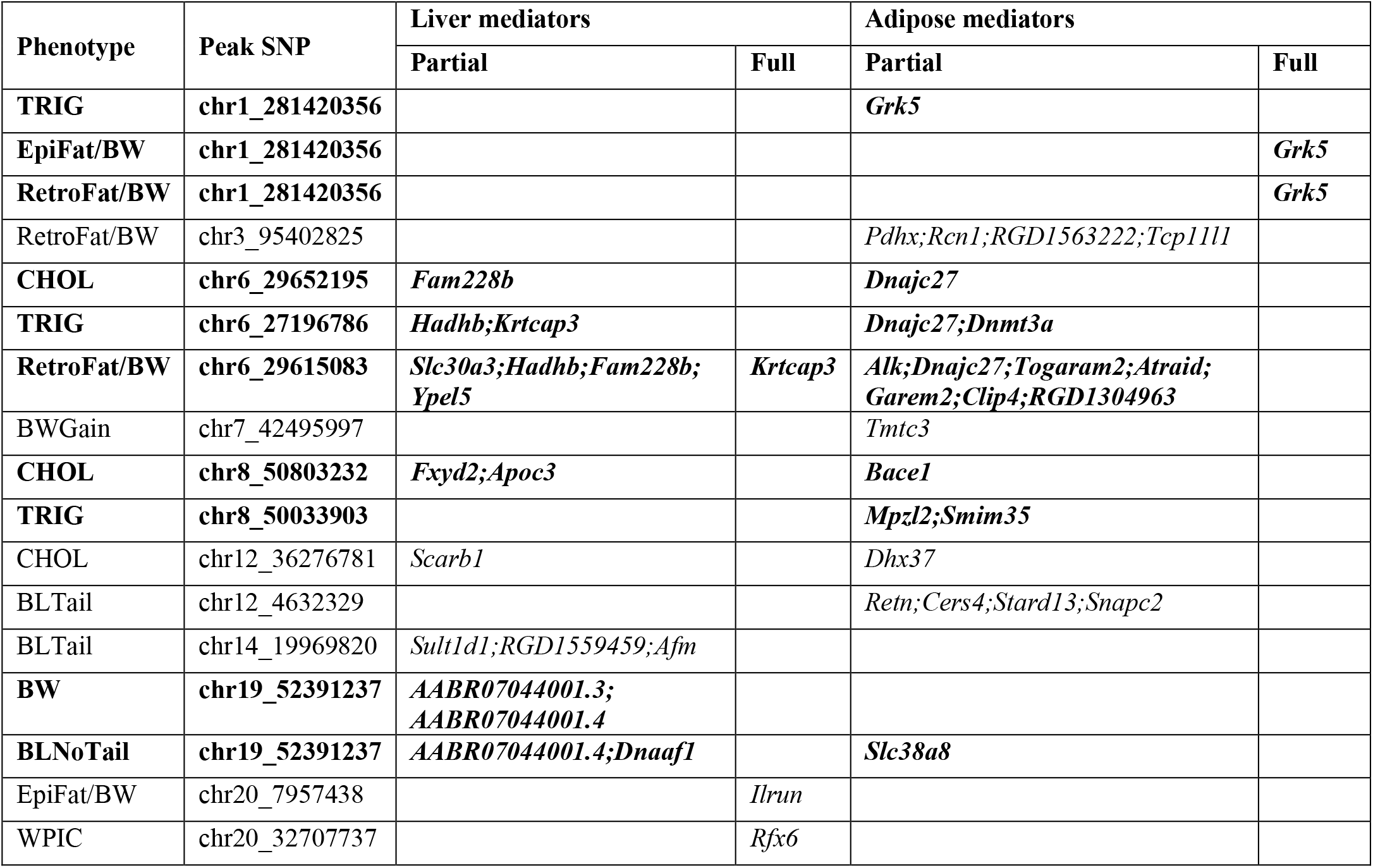
Full and Partial Mediators within each pQTL. pQTL are labelled by the peak associated SNP as in Table 1. Pleiotropic loci have been bolded.

### Full mediator: Grk5

There are 12 genes within the pQTL on Chr1:280-282 Mb for RetroFat/BW, EpiFat/BW, and TRIG, but only liver *Cacul1* and adipose *Grk5* cis-eQTLs overlap this region (**Supplementary Table 5**). Mediation analysis identified only adipose *Grk5* as a full mediator for RetroFat/BW and EpiFat/BW, and a partial mediator for TRIG (**Figure 3A-C**). The estimated haplotype effects at the pQTLs show HS founders WKY and BUF increase both fat pad weight, TRIG and *Grk5* expression, consistent with the positive correlation between *Grk5* and the phenotypes (0.11 < *R*_*p*_ < 0.16 – **Figure 3D**). Thus BUF and WKY haplotypes likely increase the expression of adipose *Grk5*, which then increases fat pad weight and TRIG levels.

Compared to control siRNA-treated cells, Day 0 (D0) undifferentiated *Grk5* shRNA-treated 3T3-L1 preadipocytes had reduced *Grk5* mRNA levels by 50% (p=0.029). Consistent with the findings from mediation analysis, there was significantly decreased adipocyte TRIG content in Day 7 (D7) differentiated *Grk5* knock-down 3T3-L1 adipocytes relative to control adipocytes (p = 0.003) (**Figure 3E**).

### Full mediator: Krtcap3

There are 123 genes within the pQTL on Chr6 for RetroFat/BW, CHOL and TRIG, of which 15 liver genes and 14 adipose genes had cis-eQTL (**Supplementary Table 7**). Of these, *Krtcap3* was the only full mediator for RetroFat/BW, and a partial mediator for TRIG (**Fig. 4A-B**). The common top SNP for these traits (chr6_29615083) is private to founder WKY, which is associated with decreased fat pad weight and increased *Krtcap3* expression. This is consistent with the negative correlation between *Krtcap3* liver expression and associated phenotypes (*R*_*p* ~_ −0.25 – **Figure 4C**), suggesting that the WKY haplotype increases the expression of liver *Krtcap3* leading to reduced accumulation of fat pad and TRIG. We have recently shown that *Krtcap3* knock-out rats exhibit increased body weight, with female rats showing increased fat pad weights and insulin sensitivity [32], thereby validating this gene. Several additional genes were identified as partial mediators, with different genes acting in adipose vs liver (**Table 2**), indicating multiple causal genes may underlie this locus.

**Figure 4 –.**
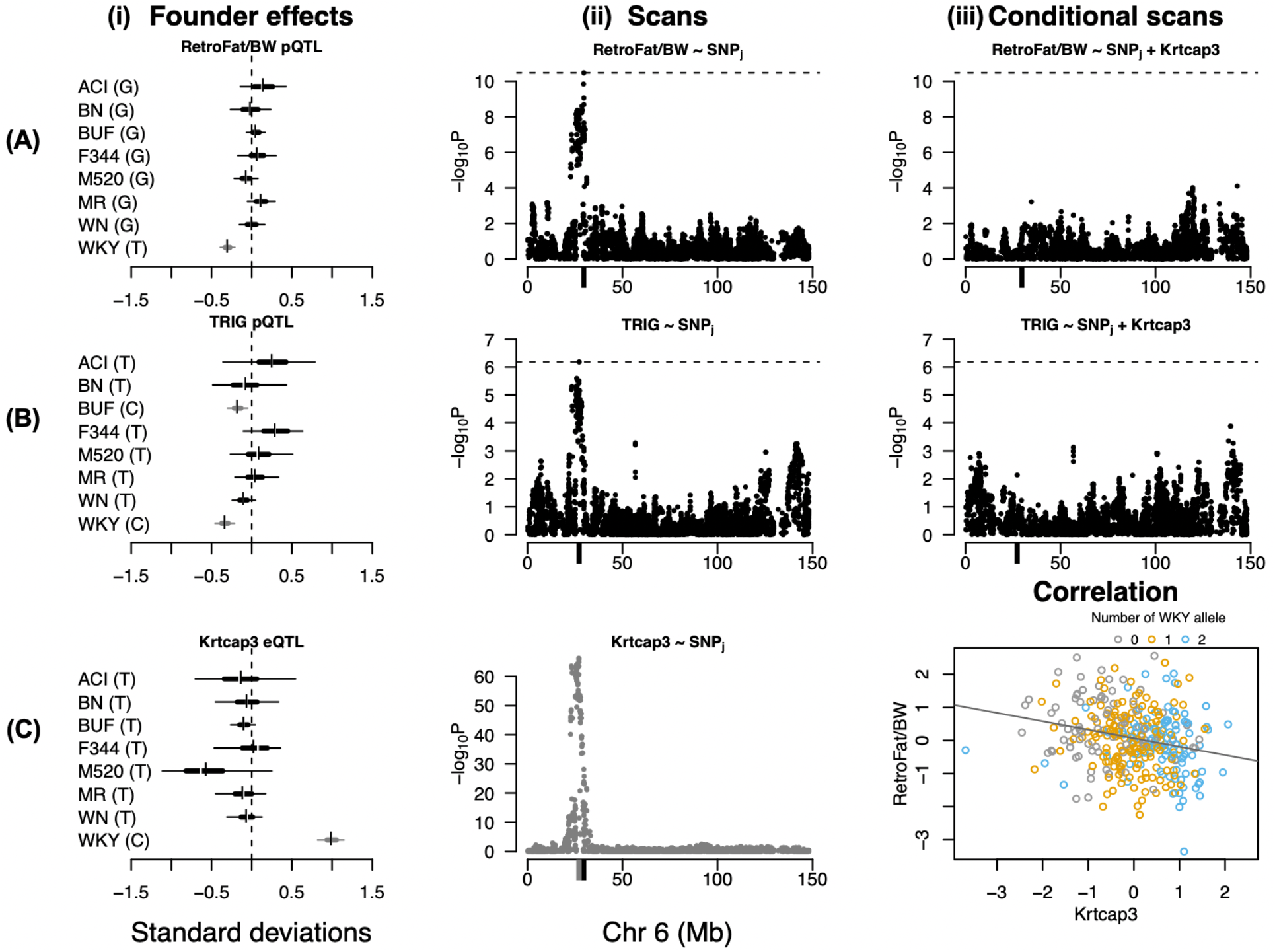
Liver expression of *Krtcap3* is full mediator for RetroFat/BW and partial mediator for Triglycerides (TRIG). See **Figure 3** for description of panels.

### Full mediator: Rfx6

There are 25 genes within the 1.4 Mb pQTL on Chr20:31.43-32.83 Mb for WPIC, of which four liver-expressed genes and three adipose-expressed genes map cis-eQTLs. Of these, liver *Rfx6* is a full mediator for WPIC. The founder haplotypes ACI and WKY are associated with decreased WPIC and *Rfx6* expression, consistent with the positive correlation between *Rfx6* and WPIC (*R*_*p*_ ~ 0.15 – **Figure 5**). These data suggest that the ACI and WKY hapolotypes decreased *Rfx6* expression leading to decreased WPIC.

**Figure 5 -.**
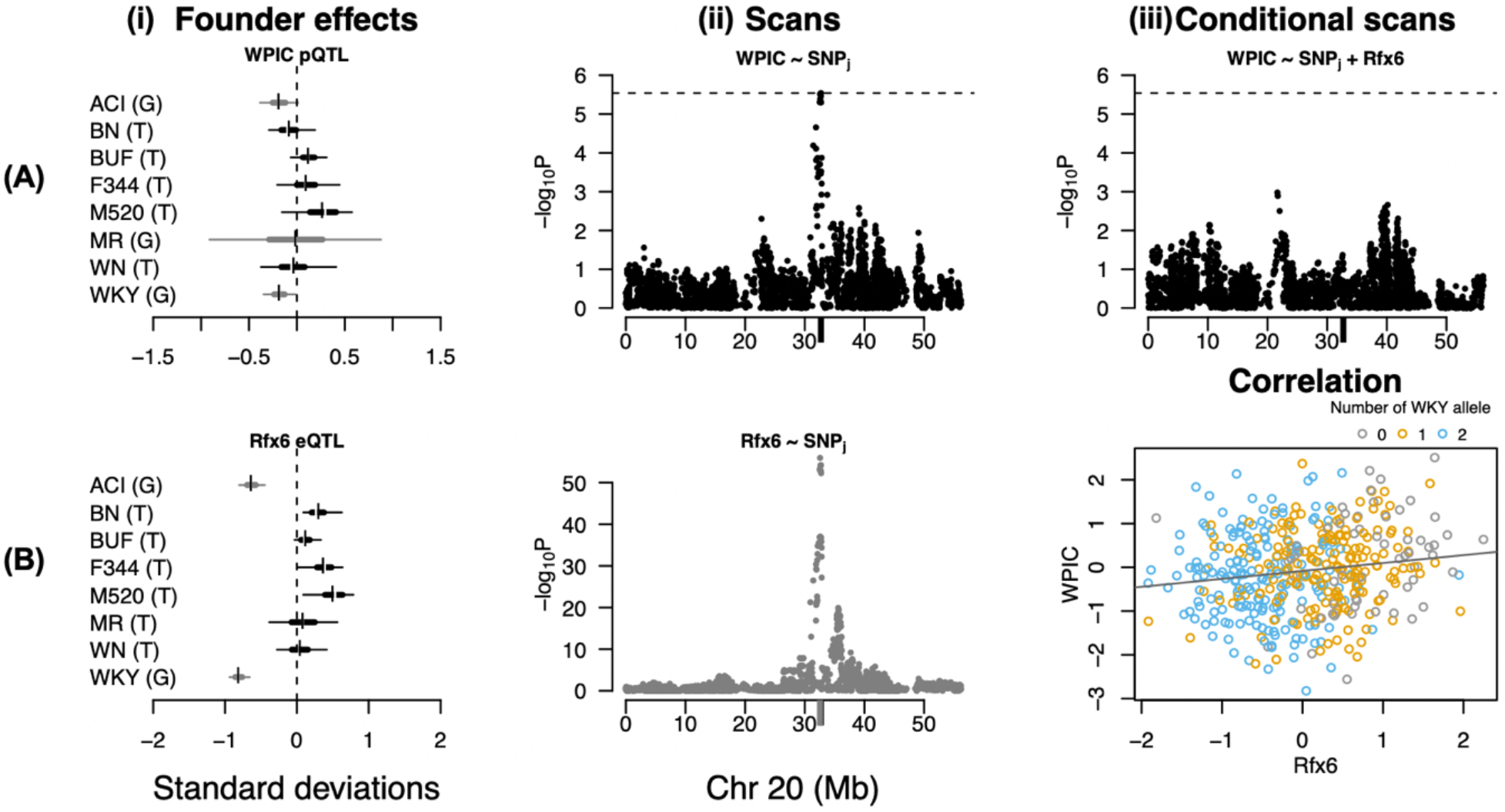
Liver expression of *Rfx6* is full mediator for Whole pancreas insulin content (WPIC). See **Figure 3** for description of panels.

### Full mediator: Ilrun

There are 27 genes within the 0.79 Mb pQTL on Chr20:7.28-8.08 Mb for EpiFat/BW, of which seven liver and ten adipose genes have cis-eQTLs. Although expressed in both tissues, only liver *Ilrun* is a full mediator for EpiFat/BW. The founder haplotype WKY is associated with both decreased EpiFat/BW and *Ilrun* liver expression. Their positive correlation (*R*_*p*_ ~ 0.17 – **Figure 6**) suggests WKY reduces *Ilrun* expression which decreases EpiFat/BW.

**Figure 6 -.**
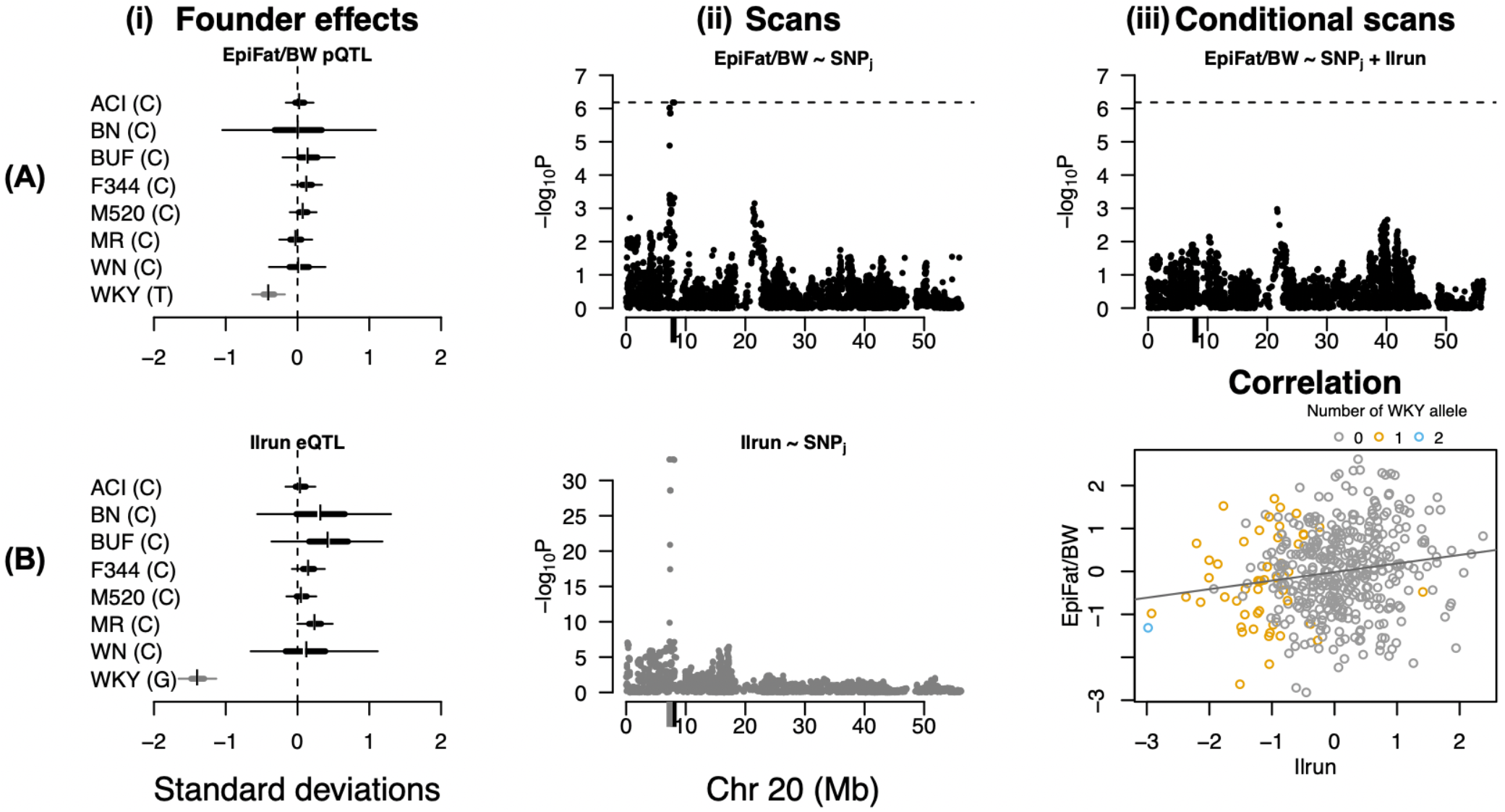
Liver expression of *Ilrun* is full mediator for EpiFat/BW. See **Figure 3** for description of panels.

## DISCUSSION

Although many loci have been identified in human GWAS for BMI and WHR [8,9], the causal genes underlying many of these loci remain unknown. In this study we addressed this challenge in outbred HS rats by collecting transcriptome data in two metabolically active tissues from the same animals for which we had in-depth metabolic phenotypes, enabling us to assess causality. We mapped 24 pQTLs for 13 metabolic traits, including five pleiotropic loci. We used mediation analysis to identify candidate causal genes, showing that multiple tissue-specific mediators underlie several loci. We identified four full mediators and verified two novel genes (*Krtcp3* and *Grk5*) experimentally. Our work demonstrates the effectiveness of imputation from LC-WGS for genotyping outbred populations, and the power of mediation analysis for identifying candidate causal genes that regulate physiological phenotypes.

In addition to identifying causal mediators, this work extends our previous work [10,11] by investigating a broader spectrum of metabolic traits including fasting lipids, glucose tolerance, insulin sensitivity and whole pancreas insulin content. We also employed both SNP-based and haplotype-based mapping; the latter of which can reveal which founder alleles have contrasting phenotypic effects and whether they have a more complex structure, potentially suggesting the presence of multiple linked causal variants [33]. In addition, we genotyped using LC-WGS followed by imputation, thereby increasing the number of markers from previous work [10]. We recommend the LC-WGS/imputation strategy as an economic way to genotype populations; it also provides data for future structural variation analysis [35].

By increasing the numbers of SNPs and animals, we replicated known adiposity loci [10] at higher significance, and mapped additional loci. We also replicated a subset of BW and fat pad weight pQTLs from a mega-analysis in HS rats [11]. In addition to 13 loci for BW and fat pad traits, we identified 3 loci for CHOL, 4 for TRIG, 2 loci for WPIC, and 4 loci for tissue weights. Surprisingly, we mapped no genome-wide significant loci for the glucose and insulin traits, despite strong heritability, suggesting these traits are highly polygenic and larger sample populations are needed. We identified full mediators underlying four loci, with partial mediators underlying several additional loci. Liver and adipose mediators often differ, demonstrating that causal genes are likely tissue-specific. We critically discuss the evidence for causality for each full mediator below.

Adipose *Grk5* is a full mediator for EpiFat/BW and RetroFat/BW on rat Chr1 and a partial mediator for TRIG. Previous work has shown that *Grk5* null mice show protection against diet-induced obesity, reduced expression of adipogenesis genes, glucose tolerance and insulin sensitivity compared with wild-type littermates [37,38]. Human GWASs have associated *Grk5* with type-2 diabetes in Chinese cohorts [39,40] and lipid levels in Europeans [7]. Our current work demonstrates that *Grk5* knock-down in 3T3L1 cells leads to decreased TRIG accumulation in adipocytes, validating a role for this gene in adiposity. Within the same pQTL there is a nonsynonymous coding variant in the gene *Prlhr* private to the BUF and WKY founders, and which was previously suggested as a candidate [10,11]. Although *Prlhr* is associated with feeding behavior [41,42], *Prlhr* is mainly expressed in brain and adrenal tissue, so our expression data could not shed light on its role as a mediator, and it is possible that both *Grk5* and *Prlhr* are causal.

Liver *Ktrcap3* is a full mediator for RetroFat/BW at the Chr6 pQTL, replicating previous work [10]. *Krtcap3* does not fall within human GWAS loci for BMI or WHR, but has been identified as a pleiotropic gene for obesity, dyslipidemia and type 2 diabetes using a multi-variate analysis [43]. Recent work by our group confirmed *Ktrcap3’*s role in adiposity using a rat *Krtcap3* knock-out; both male and female knock-out rats have increased body weight, with females showing increased adiposity and insulin sensitivity [32]. Interestingly, *Krtcap3* is unlikely the only gene acting in this locus. *Adcy3* also falls in this region and was previously nominated as a functional candidate in the rat [10,11], via a coding variant as opposed to expression levels. *Adcy3* is associated with monogenic and polygenic obesity in humans[44,45,46,47], and has been shown to act via coding variation that impacts protein folding or function [48,49], and altered expression [44]. Based on our functional validation of *Krtcap3* [32] coupled with strong support for *Adcy3* in human studies, it is likely both genes are causal for adiposity. In addition, our work identifies several partial mediators in both liver and adipose tissue, emphasizing the complexity of the Chr6 adiposity locus.

Liver *Ilrun* expression is a full mediator for EpiFat/BW on rat Chr20. *Ilrun* was previously named *LOC294154* in rats and *C6orf106* in humans. *C6orf106* associates with anthropometric traits including height [50], weight [51], BMI [52], body fat ratio [53] in humans and might drive persistent thinness versus severe obesity [54]. The mouse ortholog is implicated in lipid metabolism and hepatic lipoprotein production [55], making this a highly attractive candidate at this locus.

Liver *Rfx6* expression is a full mediator for WPIC at a separate Chr20 locus. *Rfx6* plays a role in controlling the formation of pancreatic islets and production of insulin in mice and humans [56]. Inactivation of *Rfx6* in adult beta cells impairs insulin secretion via altered glucose sensing [57]. In addition, *Rfx6* associates with monogenic diabetes of the young [58]. Together, this evidence suggests that genetic variation in *Rfx6* contributes to metabolic dysfunction and/or diabetes susceptibility.

## Conclusions

Utilizing multiple metabolic phenotypes and transcriptome data from relevant tissues from the same animals, we identify four candidate causal genes and experimentally validate two of them. This work demonstrates the power of mediation analysis for delineating genetic effects underlying multiple related phenotypes. Furthermore, it emphasizes tissue specificity of genetic effect and the potentially complex underpinnings of several metabolic loci due to multiple candidate gene drivers.

## Acknowledgements

We would like to thank Apurva Chitre for assistance in creating Porcupine plot. We would also like to thank Michael Scott for helping with STITCH. Leah C. Solberg Wood is the guarantors of this work and, as such, had full access to all the data in the study and takes responsibility for the integrity of the data and the accuracy of the data analysis.

## Author Contributions

conceived experiments (LSW, RM, WV), conducted animal experiments (LSW, KH), extracted DNA or RNA (KH, OS), RNAseq (MT, AC, GH), statistical analysis (THL, WLC, GRK), interpretation and presentation of analyzed results (THL, WV, RM, LSW), conducted 3T3L1 knock-down experiments (SKD, BM, NKS, CK), oversaw all experimental work (LSW), oversaw statistical analysis (WV, RM), wrote manuscript (THL, WV, RM, LSW). All authors approved final version of the manuscript.

## References

1. GBD 2015 Obesity Collaborators, Afshin A, Forouzanfar MH et al. Health Effects of Overweight and Obesity in 195 Countries over 25 Years. N Engl J Med 2017; 377:13–27.

2. Maes HH, Neale MC, Eaves LJ. Genetic and environmental factors in relative body weight and human adiposity. Behav Genet 1997; 27:325–351.

3. Klarin D, Damrauer SM, Cho K et al. Genetics of blood lipids among ~300,000 multi-ethnic participants of the Million Veteran Program. Nat Genet 2018; 50:1514–1523.

4. Chen J, Spracklen CN, Marenne G et al. The trans-ancestral genomic architecture of glycemic traits. Nat Genet 2021; 53:840–860.

5. Horikoshi M, Mägi R, Bunt M van de et al. Discovery and Fine-Mapping of Glycaemic and Obesity-Related Trait Loci Using High-Density Imputation. PLOS Genet 2015; 11:e1005230.

6. Vujkovic M, Keaton JM, Lynch JA et al. Discovery of 318 new risk loci for type 2 diabetes and related vascular outcomes among 1.4 million participants in a multi-ancestry meta-analysis. Nat Genet 2020; 52:680–691.

7. Yengo L, Sidorenko J, Kemper KE et al. Meta-analysis of genome-wide association studies for height and body mass index in ~700000 individuals of European ancestry. Hum Mol Genet 2018; 27:3641–3649.

8. Liu C-T, Monda KL, Taylor KC et al. Genome-Wide Association of Body Fat Distribution in African Ancestry Populations Suggests New Loci. PLOS Genet 2013; 9:e1003681.

9. Pulit SL, Stoneman C, Morris AP et al. Meta-analysis of genome-wide association studies for body fat distribution in 694 649 individuals of European ancestry. Hum Mol Genet 2019; 28:166–174.

10. Keele GR, Prokop JW, He H et al. Genetic Fine-Mapping and Identification of Candidate Genes and Variants for Adiposity Traits in Outbred Rats: Mapping Adiposity Traits in Outbred Rats. Obesity 2018; 26:213–222.

11. Chitre AS, Polesskaya O, Holl K et al. Genome wide association study in 3,173 outbred rats identifies multiple loci for body weight, adiposity, and fasting glucose. Obes Silver Spring Md 2020; 28:1964–1973.

12. Hansen C, Spuhler K. Development of the National Institutes of Health genetically heterogeneous rat stock. Alcohol Clin Exp Res 1984; 8:477–479.

13. Ramdas S, Ozel AB, Treutelaar MK et al. Extended regions of suspected mis-assembly in the rat reference genome. Sci Data 2019; 6:39.

14. Woods LCS, Mott R. Heterogeneous Stock Populations for Analysis of Complex Traits. Methods Mol Biol Clifton NJ 2017; 1488:31–44.

15. Solberg Woods LC, Palmer AA. Using Heterogeneous Stocks for Fine-Mapping Genetically Complex Traits. Methods Mol Biol Clifton NJ 2019; 2018:233–247.

16. Solberg Woods LC, Holl KL, Oreper D et al. Fine-mapping diabetes-related traits, including insulin resistance, in heterogeneous stock rats. Physiol Genomics 2012; 44:1013–1026.

17. Keele GR, Prokop JW, He H et al. Sept8/SEPTIN8 involvement in cellular structure and kidney damage is identified by genetic mapping and a novel human tubule hypoxic model. Sci Rep 2021; 11:2071.

18. Dobin A, Davis CA, Schlesinger F et al. STAR: ultrafast universal RNA-seq aligner. Bioinforma Oxf Engl 2013; 29:15–21.

19. Love MI, Huber W, Anders S. Moderated estimation of fold change and dispersion for RNA-seq data with DESeq2. Genome Biol 2014; 15:550.

20. Davies RW, Flint J, Myers S et al. Rapid genotype imputation from sequence without reference panels. Nat Genet 2016; 48:965–969.

21. Purcell S, Neale B, Todd-Brown K et al. PLINK: a tool set for whole-genome association and population-based linkage analyses. Am J Hum Genet 2007; 81:559–575.

22. Broman KW, Gatti DM, Simecek P et al. R/qtl2: Software for Mapping Quantitative Trait Loci with High-Dimensional Data and Multiparent Populations. Genetics 2019; 211:495–502.

23. Valdar W, Holmes CC, Mott R et al. Mapping in structured populations by resample model averaging. Genetics 2009; 182:1263–1277.

24. Solberg Woods LC, Holl K, Tschannen M et al. Fine-mapping a locus for glucose tolerance using heterogeneous stock rats. Physiol Genomics 2010; 41:102–108.

25. Yang J, Lee SH, Goddard ME et al. GCTA: A Tool for Genome-wide Complex Trait Analysis. Am J Hum Genet 2011; 88:76–82.

26. Zhang Z, Wang W, Valdar W. Bayesian modeling of haplotype effects in multiparent populations. Genetics 2014; 198:139–156.

27. Shabalin AA. Matrix eQTL: ultra fast eQTL analysis via large matrix operations. Bioinformatics 2012; 28:1353–1358.

28. Baron RM, Kenny DA. The moderator–mediator variable distinction in social psychological research: Conceptual, strategic, and statistical considerations. J Pers Soc Psychol 1986; 51:1173–1182.

29. Zebisch K, Voigt V, Wabitsch M et al. Protocol for effective differentiation of 3T3-L1 cells to adipocytes. Anal Biochem 2012; 425:88–90.

30. Cuffe H, Liu M, Key C-CC et al. Targeted deletion of adipocyte Abca1 impairs diet-induced obesity. Arterioscler Thromb Vasc Biol 2018; 38:733–743.

31. Key C-CC, Liu M, Kurtz CL et al. Hepatocyte ABCA1 Deletion Impairs Liver Insulin Signaling and Lipogenesis. Cell Rep 2017; 19:2116–2129.

32. Szalanczy AM, Goff E, Seshie O et al. Keratinocyte-Associated Protein 3 is a novel gene for adiposity with differential effects in males and females. 2022 2022.02.20.481201 https://www.biorxiv.org/content/10.1101/2022.02.20.481201v1 (24 February 2022, date last accessed).

33. Baud A, Hermsen R, Guryev V et al. Combined sequence-based and genetic mapping analysis of complex traits in outbred rats. Nat Genet 2013; 45:767–775.

35. Imprialou M, Kahles A, Steffen JG et al. Genomic Rearrangements in Arabidopsis Considered as Quantitative Traits. Genetics 2017; 205:1425–1441.

36. Chitre AS, Polesskaya O, Holl K et al. Genome-Wide Association Study in 3,173 Outbred Rats Identifies Multiple Loci for Body Weight, Adiposity, and Fasting Glucose. Obesity 2020; 28:1964–1973.

37. Wang F, Wang L, Shen M et al. GRK5 deficiency decreases diet-induced obesity and adipogenesis. Biochem Biophys Res Commun 2012; 421:312–317.

38. Wang L, Shen M, Wang F et al. GRK5 ablation contributes to insulin resistance. Biochem Biophys Res Commun 2012; 429:99–104.

39. Li H, Gan W, Lu L et al. A Genome-Wide Association Study Identifies GRK5 and RASGRP1 as Type 2 Diabetes Loci in Chinese Hans. Diabetes 2013; 62:291–298.

40. Xia Z, Yang T, Wang Z et al. GRK5 Intronic (CA)n Polymorphisms Associated with Type 2 Diabetes in Chinese Hainan Island. PLoS ONE 2014; 9:e90597.

41. Matta CA. Maternal treatment with teratogen causes congenital malformations in mouse embryos. Folia Morphol 1990; 38:12–18.

42. Jianjing Y, Xiaojie W, Xiaoting W et al. The Multiple Roles of XBP1 in Regulation of Glucose and Lipid Metabolism. Curr Protein Pept Sci 2017; 18:630–635.

43. Chen Y-C, Xu C, Zhang J-G et al. Multivariate analysis of genomics data to identify potential pleiotropic genes for type 2 diabetes, obesity and dyslipidemia using Meta-CCA and gene-based approach. PLOS ONE 2018; 13:e0201173.

44. Saeed S, Bonnefond A, Tamanini F et al. Loss-of-function mutations in ADCY3 cause monogenic severe obesity. Nat Genet 2018; 50:175–179.

45. Grarup N, Moltke I, Andersen MK et al. Loss-of-function variants in ADCY3 increase risk of obesity and type 2 diabetes. Nat Genet 2018; 50:172–174.

46. Nordman S, Abulaiti A, Hilding A et al. Genetic variation of the adenylyl cyclase 3 (AC3) locus and its influence on type 2 diabetes and obesity susceptibility in Swedish men. Int J Obes 2005 2008; 32:407–412.

47. Speliotes EK, Willer CJ, Berndt SI et al. Association analyses of 249,796 individuals reveal 18 new loci associated with body mass index. Nat Genet 2010; 42:937–948.

48. Stergiakouli E, Gaillard R, Tavaré JM et al. Genome-wide association study of height-adjusted BMI in childhood identifies functional variant in ADCY3. Obes Silver Spring Md 2014; 22:2252–2259.

49. Toumba M, Fanis P, Vlachakis D et al. Molecular modelling of novel ADCY3 variant predicts a molecular target for tackling obesity. Int J Mol Med 2022; 49:10.

50. Weedon MN, Lango H, Lindgren CM et al. Genome-wide association analysis identifies 20 loci that influence adult height. Nat Genet 2008; 40:575–583.

51. Cho H-W, Jin H-S, Eom Y-B. A Genome-Wide Association Study of Novel Genetic Variants Associated With Anthropometric Traits in Koreans. Front Genet 2021; 12:669215.

52. Ahmad S, Poveda A, Shungin D et al. Established BMI-associated genetic variants and their prospective associations with BMI and other cardiometabolic traits: the GLACIER Study. Int J Obes 2016; 40:1346–1352.

53. Rask-Andersen M, Karlsson T, Ek WE et al. Genome-wide association study of body fat distribution identifies adiposity loci and sex-specific genetic effects. Nat Commun 2019; 10:339.

54. Riveros-McKay F, Mistry V, Bounds R et al. Genetic architecture of human thinness compared to severe obesity. PLoS Genet 2019; 15:e1007603.

55. Bi X, Kuwano T, Lee PC et al. Ilrun,a Human Plasma Lipid GWAS Locus, Regulates Lipoprotein Metabolism in Mice. Circ Res 2020; 127:1347–1361.

56. Smith SB, Qu H-Q, Taleb N et al. Rfx6 directs islet formation and insulin production in mice and humans. Nature 2010; 463:775–780.

57. Piccand J, Strasser P, Hodson DJ et al. Rfx6 Maintains the Functional Identity of Adult Pancreatic β Cells. Cell Rep 2014; 9:2219–2232.

58. Patel KA, Kettunen J, Laakso M et al. Heterozygous RFX6 protein truncating variants are associated with MODY with reduced penetrance. Nat Commun 2017; 8:888.

