## Supplementary figures and images for "Genetic mapping of multiple metabolic traits identifies novel genes for adiposity, lipids and insulin secretory capacity in outbred rats"

### Supplementary Figure 1

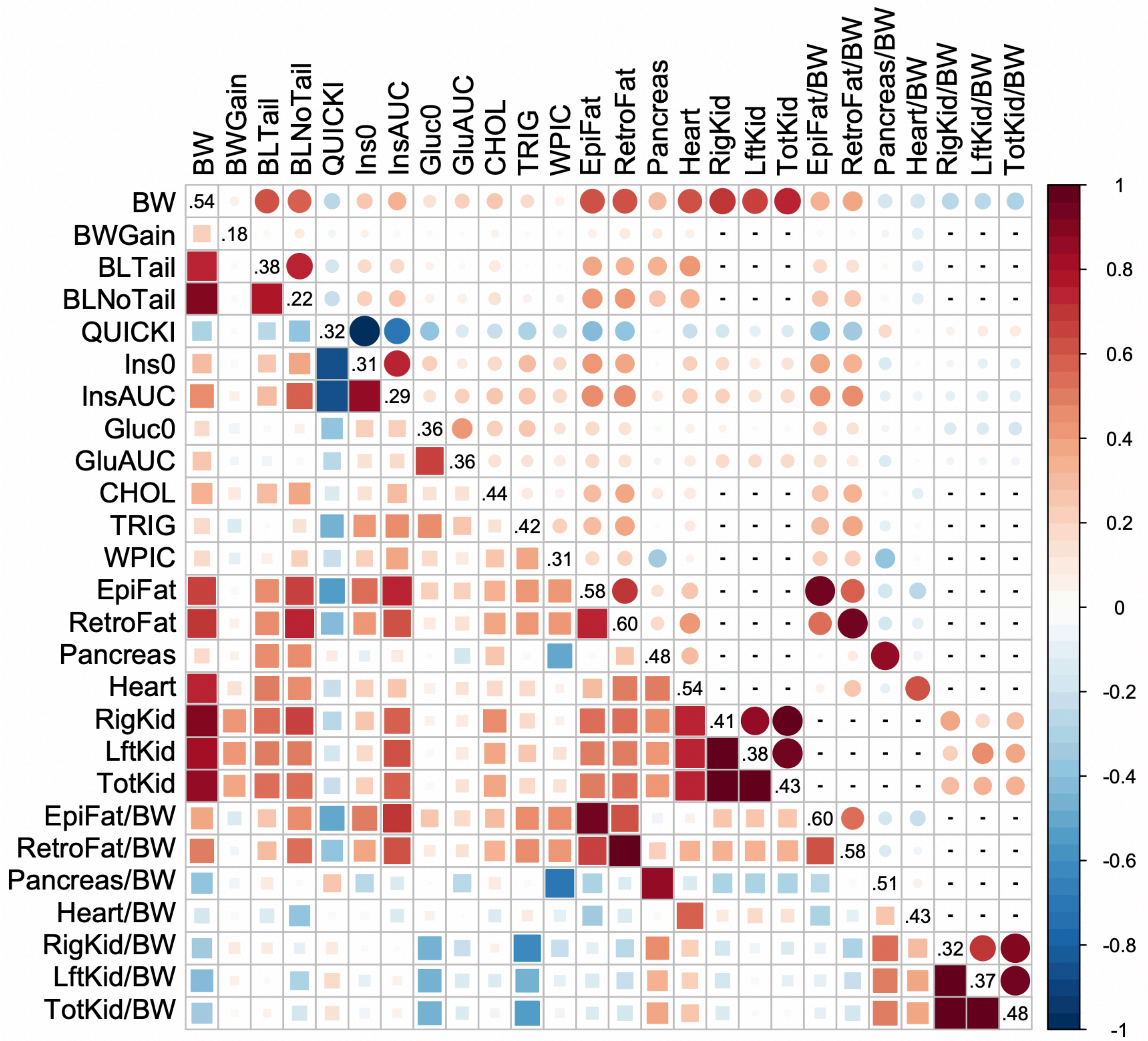

### Supplementary Figure 2

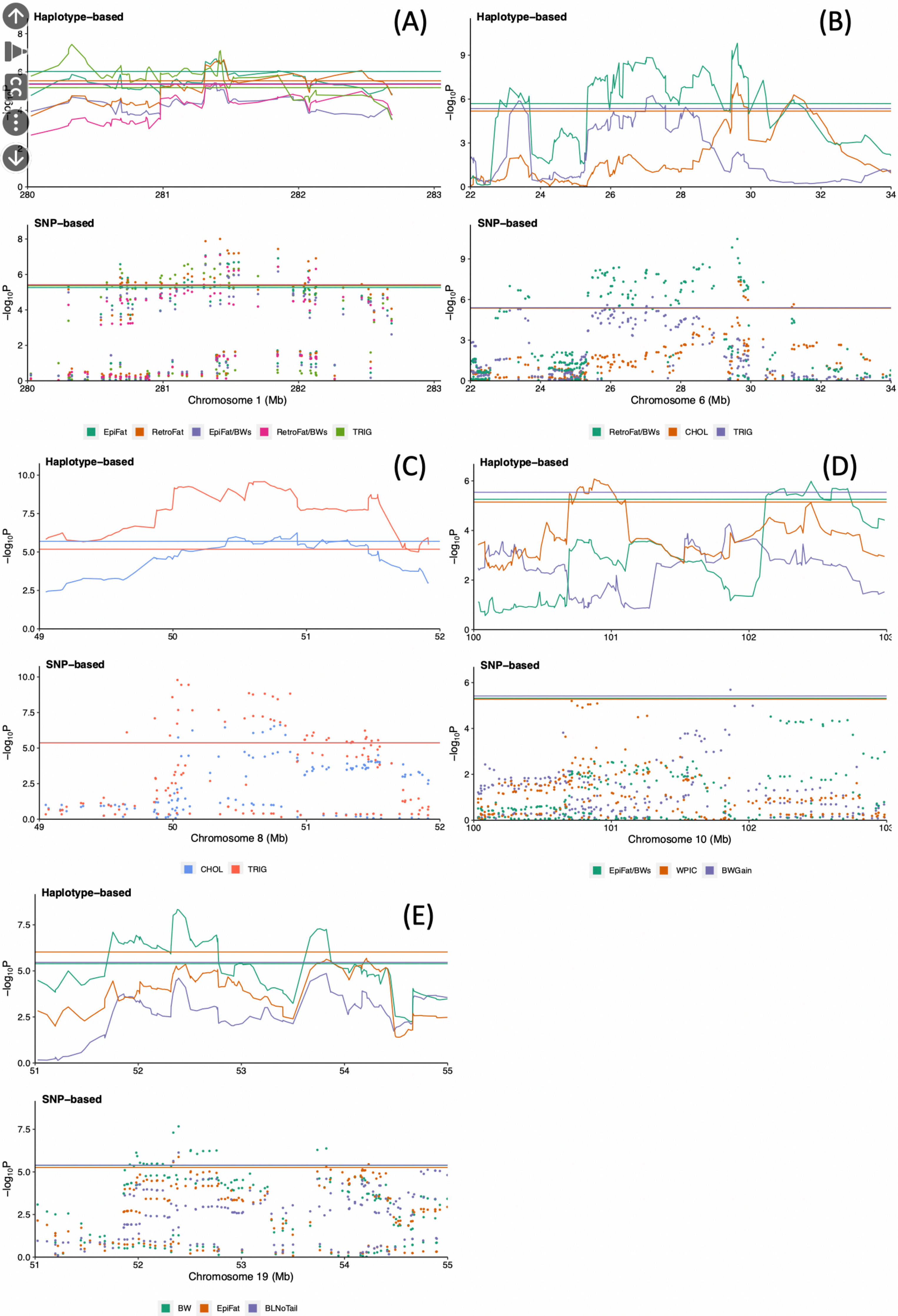
