## Supplementary Figure 3 for "Genetic mapping of multiple metabolic traits identifies novel genes for adiposity, lipids and insulin secretory capacity in outbred rats"

**Supplementary. Figure 3 -** Manhattan plots for physiological traits using (A) haplotype-based or (B) SNP-based mixed models. The x-axis shows genomic position and y-axis the significance as negative log 10 of the p-value of the test of association. Thresholds are shown as red horizontal lines and indicate the genome-wide 10% thresholds as determined by permutation in haplotype and bootstraps in SNP-based method.

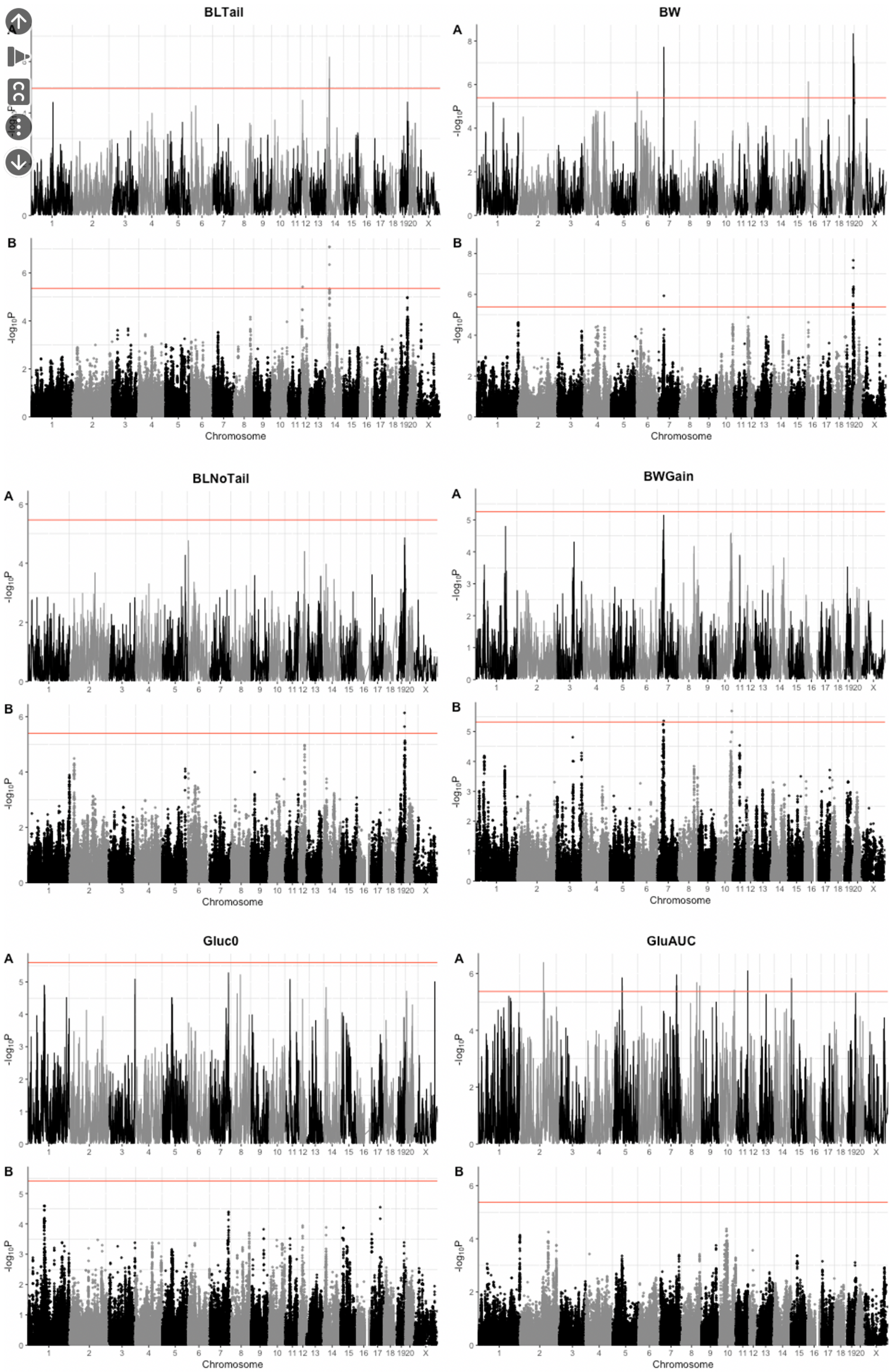
BLTail
BW
BLNoTail
BWGain
Gluc0
GluAUC
A
B
-log10P
Chromosome
1 2 3 4 5 6 7 8 9 10 11 12 13 14 15 16 17 18 19 20 X

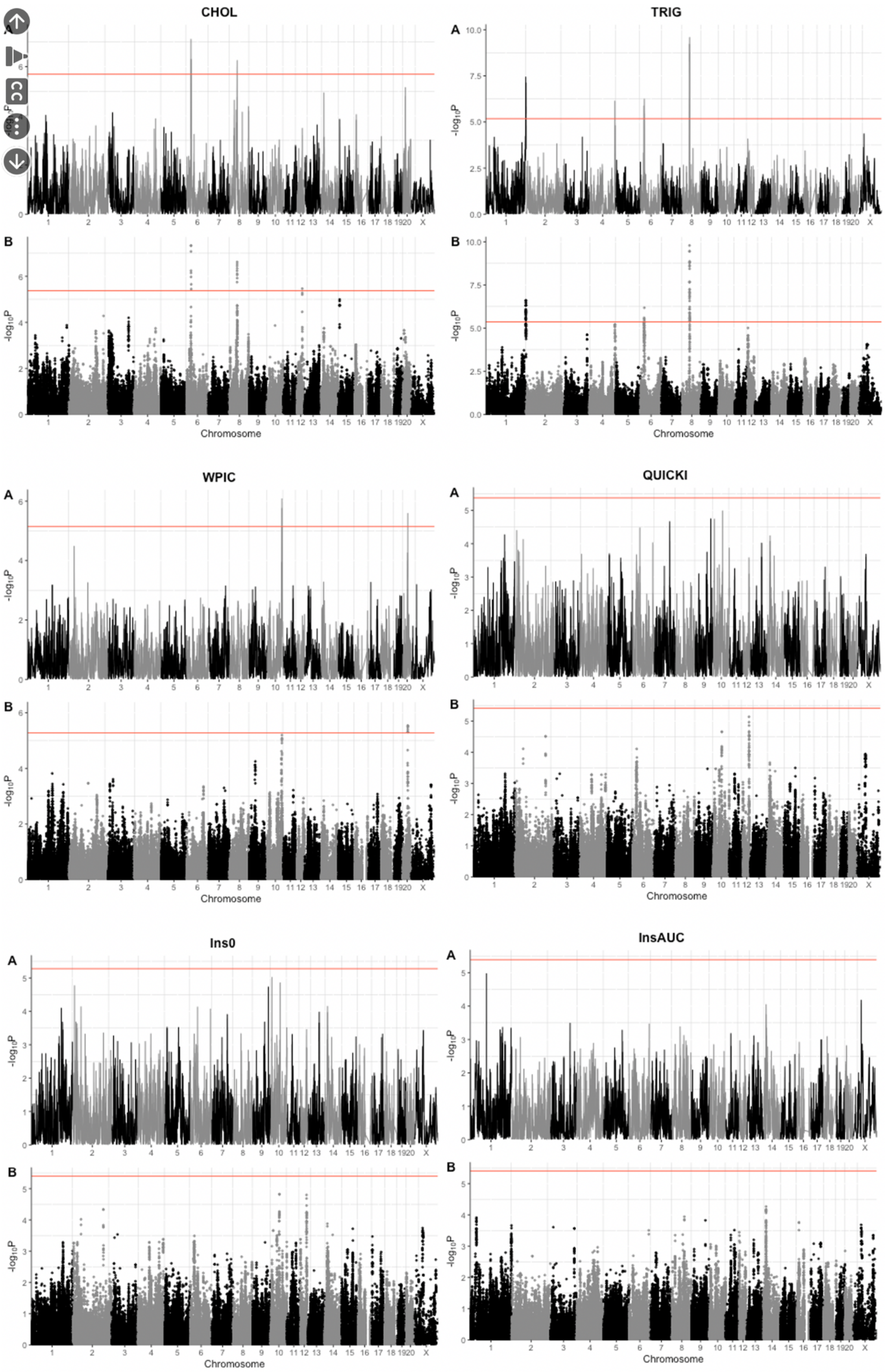
CHOL
TRIG
WPIC
QUICKI
Ins0
InsAUC
A
B
-log10P
Chromosome
1 2 3 4 5 6 7 8 9 10 11 12 13 14 15 16 17 18 1920 X

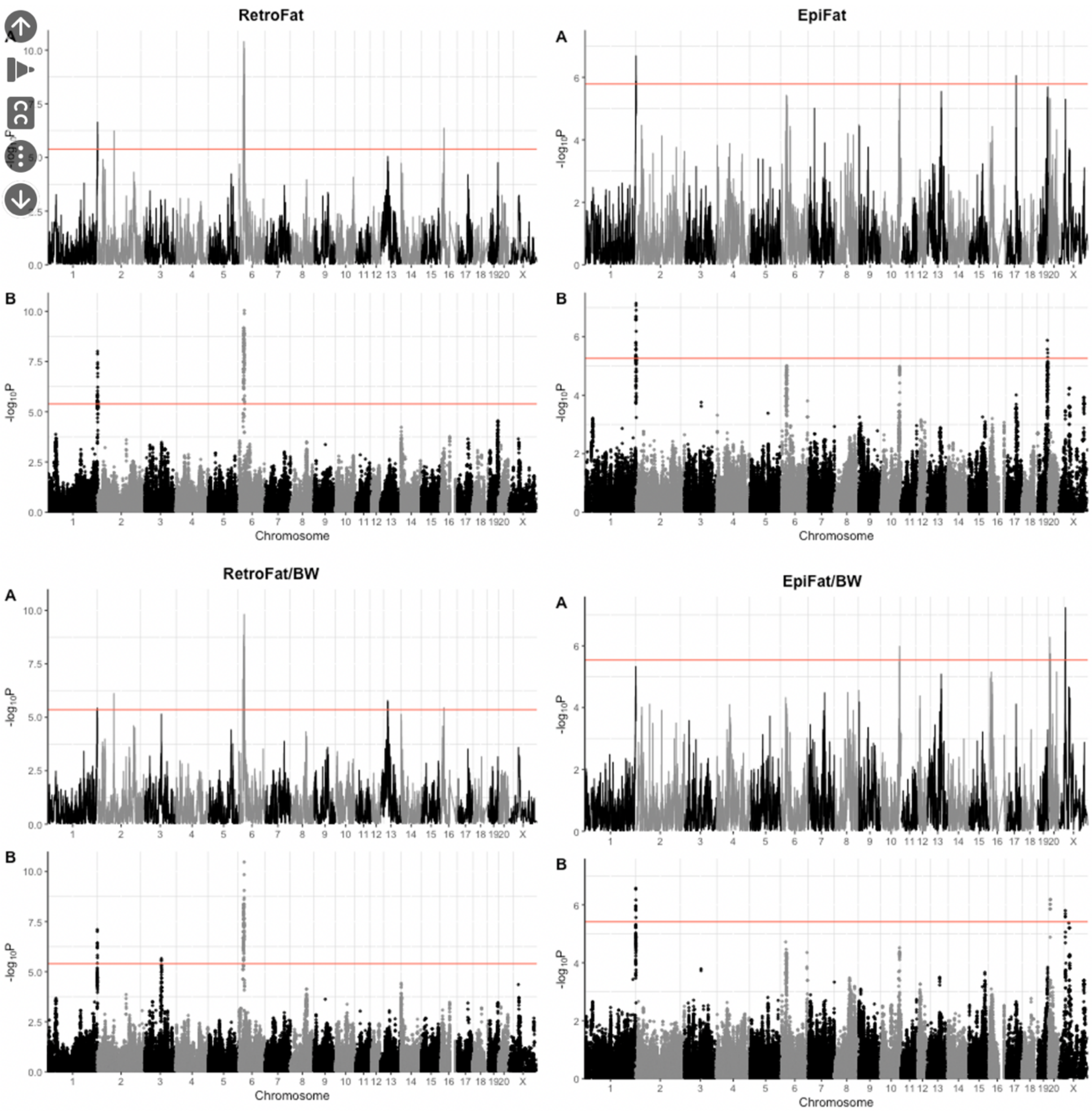
RetroFat
EpiFat
RetroFat/BW
EpiFat/BW
A
B
-log10P
Chromosome
1 2 3 4 5 6 7 8 9 10 11 12 13 14 15 16 17 18 19 20 X

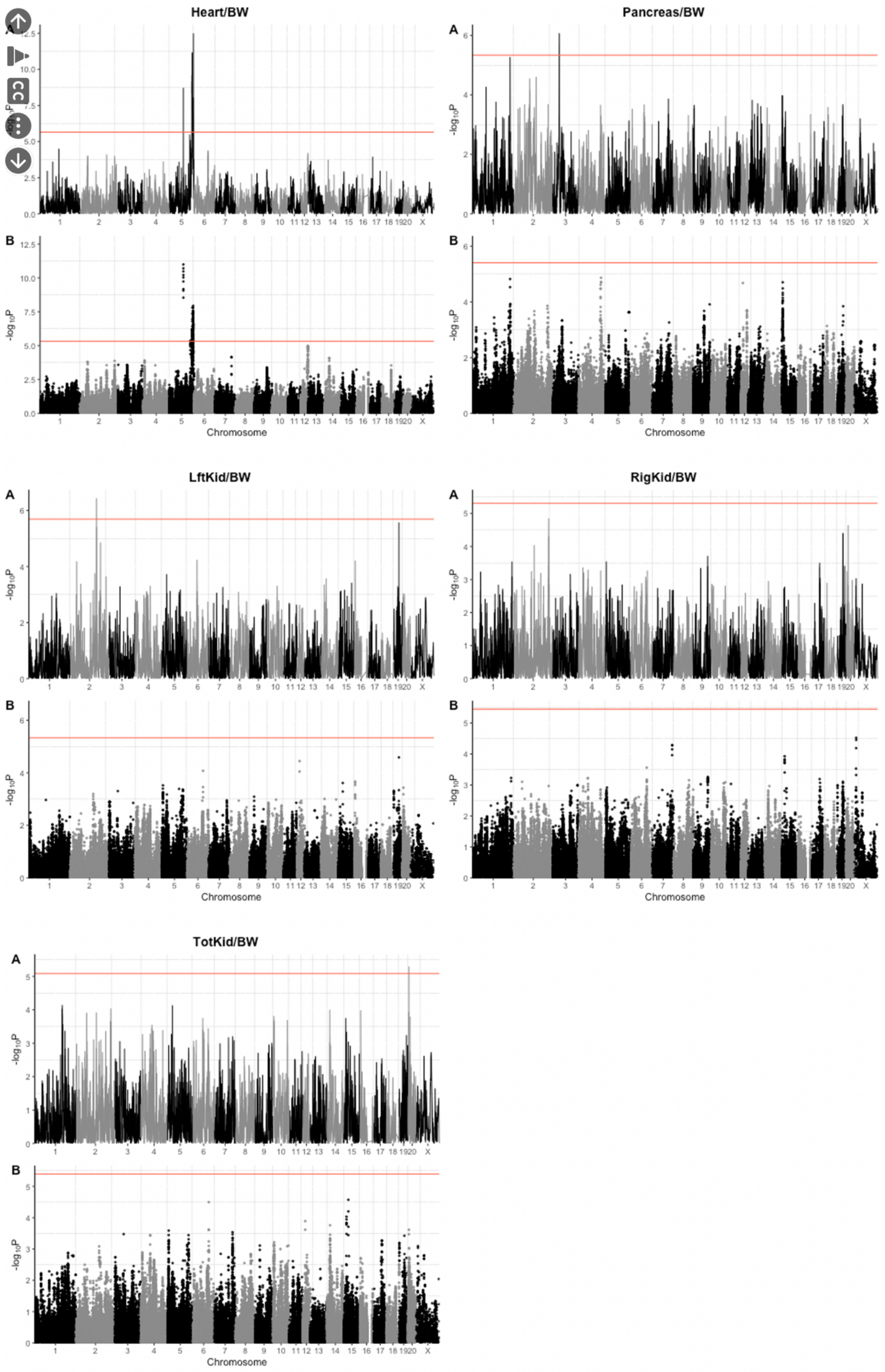
Heart/BW
Pancreas/BW
LftKid/BW
RigKid/BW
TotKid/BW
A
B
$-\log_{10}P$
Chromosome
1 2 3 4 5 6 7 8 9 10 11 12 13 14 15 16 17 18 19 20 X
